# Virtual staining from bright-field microscopy for label-free quantitative analysis of plant cell structures

**DOI:** 10.1101/2024.10.22.619768

**Authors:** Manami Ichita, Haruna Yamamichi, Takumi Higaki

**Affiliations:** Graduate School of Science and Technology, Kumamoto University, 2-39-1 Kurokami, Chuo-Ku, Kumamoto, 860-8555, Japan; Faculty of Science, Kumamoto University, 2-39-1 Kurokami, Chuo-Ku, Kumamoto, 860-8555, Japan; International Research Organization for Advanced Science and Technology, Kumamoto University, 2-39-1 Kurokami, Chuo-Ku, Kumamoto, 860-8555, Japan

**Author notes:** Correspondence should be addressed to Takumi Higaki, Graduate School of Science and Technology, Kumamoto University, 2-39-1 Kurokami, Chuo-Ku, Kumamoto, 860-8555, Japan.

**Keywords:** deep learning, pavement cells, quantitative cellular analysis, tobacco BY-2 cells, virtual staining

## Abstract

The applicability of a deep learning model for the virtual staining of plant cell structures using bright-field microscopy was investigated. The training dataset consisted of microscopy images of tobacco BY-2 cells with the plasma membrane stained with the fluorescent dye PlasMem Bright Green and the cell nucleus labeled with Histone-red fluorescent protein. The trained models successfully detected the expansion of cell nuclei upon aphidicolin treatment and a decrease in the cell aspect ratio upon propyzamide treatment, demonstrating its utility in cell morphometry. The model also accurately documented the shape of Arabidopsis pavement cells in both wild type and the *bpp125* triple mutant, which has an altered pavement cell phenotype. Metrics such as cell area, circularity, and solidity obtained from virtual staining analyses were highly correlated with those obtained by manual measurements of cell features from microscopy images. Furthermore, the versatility of virtual staining was highlighted by its application to track chloroplast movement in *Egeria densa*. The method was also effective for classifying live and dead BY-2 cells using texture-based machine learning, suggesting that virtual staining can be applied beyond typical segmentation tasks. Although this method still has some limitations, its non-invasive nature and efficiency make it highly suitable for label-free, dynamic, and high-throughput analyses in quantitative plant cell biology.

## Introduction

Determination of the shape of plant cells, which are generally immotile, is crucial because it impacts the morphogenesis of plant tissues and organs (Kierzkowski and Routier-Kierzkowska 2019; Kikukawa et al. 2023). The shape of plant cells is mainly determined by the orientation of the plane of cell division and the direction of cell elongation, processes largely regulated by the cytoskeleton and cell wall (Cosgrove 2005; Smith and Oppenheimer 2005). The analysis of molecular regulators associated with the cytoskeleton and cell wall requires an accurate and quantitative evaluation of plant cell shape in gene mutants and in samples treated with relevant inhibitors (Wolny et al. 2020; Kikukawa et al. 2021a, b). To quantitatively evaluate plant cell morphology, it is common to first obtain fluorescent images of the specimens with fluorescently labeled plasma membranes or cell walls using confocal microscopy. Subsequently, the cell regions are determined, i.e., segmented, and then the cell shape is quantitatively evaluated based on morphological metrics such as cell circularity (Kikukawa et al. 2021a). While recent advances in image analysis techniques have enabled automated and high-precision cell segmentation (Legland et al. 2016; Wolny et al. 2020; Kikukawa et al. 2021a), acquiring fluorescent images of cell contours can sometimes be a bottleneck. Undoubtedly, fluorescence imaging has dramatically propelled advancements in life sciences. However, in fluorescence imaging, phototoxicity issues are unavoidable in principle. Despite remarkable progress in fluorescent probe development (Wang et al. 2017; Hirano et al. 2022), image quality often deteriorates because of photobleaching, particularly during time-lapse observations. Moreover, potential adverse effects of invasion of the fluorescent dyes used for staining plasma membranes or cell walls should be considered. Even in the case of fluorescent proteins, adverse effects of genetic transformation cannot be dismissed. Of course, this is not applicable to biological materials where genetic transformation is challenging. With this issue in mind, we have been developing imaging analysis systems to complement existing fluorescence imaging techniques for plant cell morphology analysis. For instance, to visualize and track the morphology of leaf pavement cells, we developed a method of applying metal ink to the leaf surfaces, capturing images with a metallographic microscope, and using deep learning to highlight and segment cell contours (Kikukawa et al. 2021b). Compared with fluorescence imaging, this deep learning-powered metallographic imaging method is highly accurate, cost-effective, and applicable to non-transformable plants (Kikukawa et al. 2021b). However, it still requires the application of metal ink, the adverse effects of which cannot be completely eliminated (Kikukawa et al. 2021b).

In the field of biomedical and pharmaceutical research, there is an emerging technique that uses deep learning to convert transmitted bright-field microscopy images into synthetic fluorescence images, allowing for label-free and specific visualization of the cell structures in animal cells (Christiansen et al. 2018; Ounkomol et al. 2018; Wieslander et al. 2021; Cross-Zamirski et al. 2022). This approach, known as *in silico* labeling (Christiansen et al. 2018), label-free prediction (Ounkomol et al. 2018), or virtual staining (Rivenson et al. 2019) (hereafter referred to as virtual staining), complements physical staining, including fluorescence imaging, and has the potential to significantly advance research in biomedical and pharmaceutical sciences. This technique has potential applications not only for the segmentation and morphometric analysis of plant cells, but also for broader tasks, such as automatic recognition of cell states, which could enable versatile analyses in plant cell biology. However, the utility of virtual staining in plant cells has remained unclear.

With an appropriate training image dataset, virtual staining was shown to be feasible for tobacco BY-2 cells, enabling both multiple staining and morphometric analyses. To further assess the versatility of this virtual staining model, we applied it to the morphometric analysis of *Arabidopsis thaliana* pavement cells, demonstrating that it also has applications in high-precision cell morphometry and in phenotypic analyses of genetic mutants. Additionally, we explored the applicability of virtual staining for tracking chloroplast movement in *Egeria densa* and for the classification of live and dead BY-2 cells through texture-based machine learning. Our results show that virtual staining can support morphometric analysis, while also offering advantages for non-segmentation-based and dynamic cellular analyses in plant cell research.

## Methods

### Plant materials and fluorescent labeling

Tobacco BY-2 (*Nicotiana tabacum* L. cv. Bright Yellow 2) cells were diluted 95-fold with modified Linsmaier and Skoog medium supplemented with 2,4-dichlorophenoxyacetic acid at weekly intervals (Kumagai-Sano et al. 2006). The cell suspensions were agitated on a rotary shaker at 130 rpm at 27°C in the dark. To visualize the cell nuclei, a transgenic BY-2 cell line stably expressing red fluorescent protein-tagged Histone H2B (Histone-RFP) under the control of the cauliflower mosaic virus (CaMV) 35S promoter was used (Maeda and Higaki 2021; Okubo-Kurihara et al. 2022). To visualize the microtubules and actin microfilaments, a transgenic BY-2 cell line stably expressing yellow fluorescent protein-tagged β-tubulin (YFP-β-tubulin) and Lifeact-mCherry, both under the control of the CaMV 35S promoter, was used (Yasuhara and Kitamoto 2014). To visualize the plasma membranes, 0.5 μL of the fluorescent dye PlasMem Bright Green (Dojindo, Kumamoto, Japan) (Liu et al. 2022) was added to 100 μL of cell suspension, and then the cells were observed immediately.

Seeds of wild-type *A. thaliana* (Col-0), the *bpp125* triple mutant (Wong et al. 2019; Yoshida et al. 2022), and the transgenic line expressing green fluorescent protein-tagged plasma membrane intrinsic protein 2a (GFP-PIP2a), which is a plasma membrane marker (Cutler et al. 2000; Kikukawa et al. 2021a, b), were sown in culture soil (Jiffy-7; Sakata Seed Corp., Kanagawa, Japan) in plastic pots and grown in a controlled-environment chamber (LH-241PFP-S; NK system, Osaka, Japan) at 23.5°C under a 16-h light/8-h dark photoperiod. To observe the leaf epidermis, the abaxial epidermis of the mature true leaf was peeled off and mounted on a slide with sterilized half-strength Murashige and Skoog liquid medium (Higaki et al. 2010).

*E. densa* was obtained from a commercial supplier (1-157-0711; Kenis Ltd., Osaka, Japan) for use as an experimental sample. To observe chloroplast movement, a detached leaflet was mounted in sterilized water, and the adaxial surface was observed.

### Inhibitor treatments

To pharmacologically induce cell nuclei deformation and morphological changes in BY-2 cells, 0-day-old cell suspension cultures were treated with DMSO, 5 mg/L aphidicolin, a DNA polymerase inhibitor, or 10 μM propyzamide, a tubulin polymerization inhibitor. The cells were cultured for 3 days before observation.

To obtain the training images of chloroplasts in *E. densa*, 100 mM 2,3-butanedion monoxime (BDM), a myosin ATPase inhibitor (Higaki et al. 2006), was applied to prevent chloroplast movement during the interval between capturing bright-field images and capturing chloroplast autofluorescence images.

### Microscopy

Transmitted bright-field images of BY-2 cells and *A. thaliana* were captured with a microscope (IX-70; Olympus, Tokyo, Japan) equipped with an objective lens (UPlanXApo ×20; Olympus; NA = 0.80) and a halogen lamp house (U-LH100L-3; Olympus) for bright-field observation. The exposure time was set to 30 ms, with a spatial resolution of 0.56 μm per pixel. For bright-field image acquisition, the focus was manually adjusted to a point where the cell contour brightness was reduced. For confocal imaging, a spinning disk confocal scanning unit (CSU-W1; Yokogawa, Tokyo, Japan), a laser illumination homogenization unit (Uniformizer; Yokogawa), and a complementary metal-oxide semiconductor camera (Zyla; Andor, Belfast, United Kingdom) were used. The fluorescent dye PlasMem Bright Green as well as GFP and YFP were excited at 488 nm, with emission wavelengths of 510–550 nm. Both RFP and mCherry were excited at 561 nm, with emission wavelengths of 624–640 nm. To visualize the vacuolar lumen in BY-2 cells, the cells were stained with 10 μM BCECF, a fluorescent dye, and then cultured for 1 hour (Kutsuna and Hasezawa 2002). After staining, the cells were imaged using a confocal microscope with an excitation wavelength of 488 nm and emission wavelengths of 510–550 nm.

*E. densa* images were similarly captured with the same confocal microscope system using a UPlanApo ×40 objective lens (NA = 0.85), an exposure time of 10 ms, and a spatial resolution of 0.28 μm per pixel. Chloroplast autofluorescence was excited at 561 nm, with emission detected using a 575 nm interference filter (575IF).

### Virtual staining

Virtual staining was performed with the deep learning-based 2-D segmentation function of the image analysis software AIVIA version 9.8.1 (DRVision, Bellevue, WA, United States) (Kikukawa et al. 2021b). The segmentation model used in this study is based on the residual channel attention (RCA)-UNet architecture, which builds upon the UNet architecture (Ronneberger et al. 2015) by replacing the convolution blocks with RCA blocks (Zhang et al. 2018). This RCA-UNet architecture enhances the model’s ability to focus on informative features at each resolution level, offering improved performance over the original UNet (Zhang et al. 2018). The training parameters followed the default settings, which are detailed in the AIVIA Wiki. A complete list and explanation of these hyperparameters can be found at the following link: https://aivia-software.atlassian.net/wiki/spaces/AW/pages/1797652487/Deep+Learning+Hyperparameters+Settings.

For virtual staining of the plasma membrane and cell nuclei, we used 150 bright-field images of BY-2 cells as input examples, along with corresponding confocal images of PlasMem Bright Green-labeled plasma membranes or Histone-RFP-labeled cell nuclei in the same field of view as reference fluorescence images. To evaluate the accuracy of the virtual staining models, we tested them on a separate dataset of 43 images that were not included in the training process. For virtual staining of vacuoles, 150 bright-field images of BY-2 cells served as input examples, with corresponding confocal images of the BCECF-labeled vacuolar lumen as reference fluorescence images in the same field of view. We assessed the accuracy of the vacuole virtual staining model using a test dataset of 25 images that were excluded from the training dataset. For the virtual staining of chloroplasts in *E. densa*, we used 109 images for training and 10 images for testing. These training and test images are publicly available on figshare under the CC BY 4.0 license. For access and detailed URLs, please refer to the Data Availability section.

To quantitatively evaluate the accuracy of virtual staining, colocalization was quantified using the threshold overlap score (TOS) with logarithmic rescaling. The TOS metric was analyzed with the top 1 percentile using the ImageJ plugin EzColocalization (Stauffer et al. 2018). The TOS provides a measure of colocalization that scales logarithmically rather than linearly, where a value of −1 indicates complete anticolocalization, 0 represents non-colocalization, and 1 corresponds to complete colocalization.

### Segmentation and measurements of morphological metrics

Image analysis for segmentation and morphometric measurements was conducted using ImageJ (Schneider et al. 2012). To investigate the effects of aphidicolin or propyzamide on the cell nuclei area and cell aspect ratio, the cell nuclei regions and the regions enclosed by the plasma membrane in virtual staining images were segmented using ImageJ’s Otsu method, and the area and aspect ratio were measured using the Analyze Particles function in ImageJ. For morphometric analysis of pavement cells, manual segmentation based on bright-field images was performed using ImageJ’s Freehand tool. Semi-automated segmentation based on virtual staining images was carried out using the ImageJ plugin, Morphological Segmentation (Legland et al. 2016). The tolerance parameter, which controls the sensitivity of segmentation to intensity differences, was manually adjusted within a range of 1,200 to 14,000 based on visual inspection to optimize the segmentation for each image set. This parameter was fine-tuned for each dataset to ensure accurate segmentation of cell boundaries while minimizing over-segmentation or under-segmentation. The area, circularity, and solidity of these cells were measured. Circularity and solidity are major indicators in the evaluation of pavement cell morphology (Higaki et al. 2017; Kikukawa et al. 2021a). Circularity was defined as 4πSL^–2^, where S and L represent the cell area and perimeter, respectively. The highest value of 1 indicates a perfect circle, whereas the lowest value of 0 indicates a highly complex shape (Higaki et al. 2017; Kikukawa et al. 2021a). Solidity, which is the ratio of cell area to the area of the convex hull of the cell, was used as an index of cell interdigitation. It reaches a maximum value of 1 when there are no waves in the lateral cell wall and approaches a minimum value of 0 when the wave shape of the lateral cell wall is more pronounced (Higaki et al. 2017; Kikukawa et al. 2021a).

### Measurements of chloroplast movement

To manually measure the speed of chloroplast movement, the centroids of chloroplasts were manually identified using the Point tool in ImageJ based on visual inspection of bright-field images. For automated measurement of the speed of chloroplast movement from virtually stained images, the chloroplasts were segmented using Otsu’s thresholding method in ImageJ, and the XY coordinates of the centroids were obtained. In both manual and automated approaches, the chloroplast movement speed between frames was calculated from the XY coordinates of the chloroplast centroids.

### Machine learning classification of living and dead cells

To automatically classify living and dead cells from bright-field and virtually stained images, 0-day-old tobacco BY-2 cells were dispensed into a 96-well plate with 100 µL per well. Cells were treated with either 0.1% DMSO (control) or 10 µM enoxolone (Selleck, Kanagawa, Japan), followed by 24 hours of culture in the dark at 27°C with shaking. After incubation, bright-field images of the cells in the wells were automatically acquired using the imaging system (BioTek Cytation1, Agilent, CA, USA). From these images, 1,200 live cells from the DMSO control and 1,200 dead cells were manually cropped using ImageJ. Of these, 900 images per condition were used to train a random forest classification model, while the remaining 300 images per condition were reserved as test images for model evaluation. For feature extraction, the ImageJ plug-in LPX Features was used to calculate gray level co-occurrence matrix (GLCM) features (Haralick 1979) (https://lpixel.net/en/products/lpixel-imagej-plugins/). The classification model was trained using these features with the R package randomForest (https://cran.r-project.org/web/packages/randomForest/index.html), utilizing the default parameters (Yoshida et al. 2022).

## Results

To train a deep learning model for virtual staining of plant cell contours and intracellular structures, we collected confocal microscopy images of tobacco BY-2 cells with the plasma membrane stained with the fluorescent dye PlasMem Bright Green (Liu et al. 2022) and the cell nucleus labeled with Histone-RFP (Maeda and Higaki 2021; Okubo-Kurihara et al. 2022), along with bright-field images from the same field of view that had slightly defocused contours to emphasize shadows (Fig. 1a). We utilized a commercially available pre-trained deep learning model for 2D image segmentation in AIVIA software (Kikukawa et al. 2021b), which was trained in two separate sessions using various combinations of the two types of bright-field images and corresponding fluorescence images. The model employs UNet architecture (Ronneberger et al. 2015), where conventional convolutional blocks are replaced with RCA blocks (Zhang et al. 2018) to enhance feature extraction capabilities. Each training image set consisted of 150 pairs of bright-field and fluorescence images. The trained model successfully outputted images that corresponded well with the spatial distribution patterns of the PlasMem Bright Green or Histone-RFP (Fig. 1b). To quantitatively assess virtual staining accuracy, we used the colocalization metric, TOS (Stauffer et al. 2018; Hotta et al. 2022), across a test set of 43 pairs of bright-field and fluorescence images not included in the training dataset (Table 1). Measurements of TOS revealed that the virtually stained plasma membranes exhibited low colocalization with Histone-RFP (0.266 ± 0.249) but high colocalization PlasMem Bright Green (0.866 ± 0.0481). Conversely, the virtually stained cell nuclei showed high colocalization with Histone-RFP (0.798 ± 0.0760) and low colocalization with PlasMem Bright Green (0.465 ± 0.282) (Table 1). These results confirm that the virtual staining method accurately replicates the spatial distribution of each targeted structure with appropriate selectivity.

**Fig. 1.**
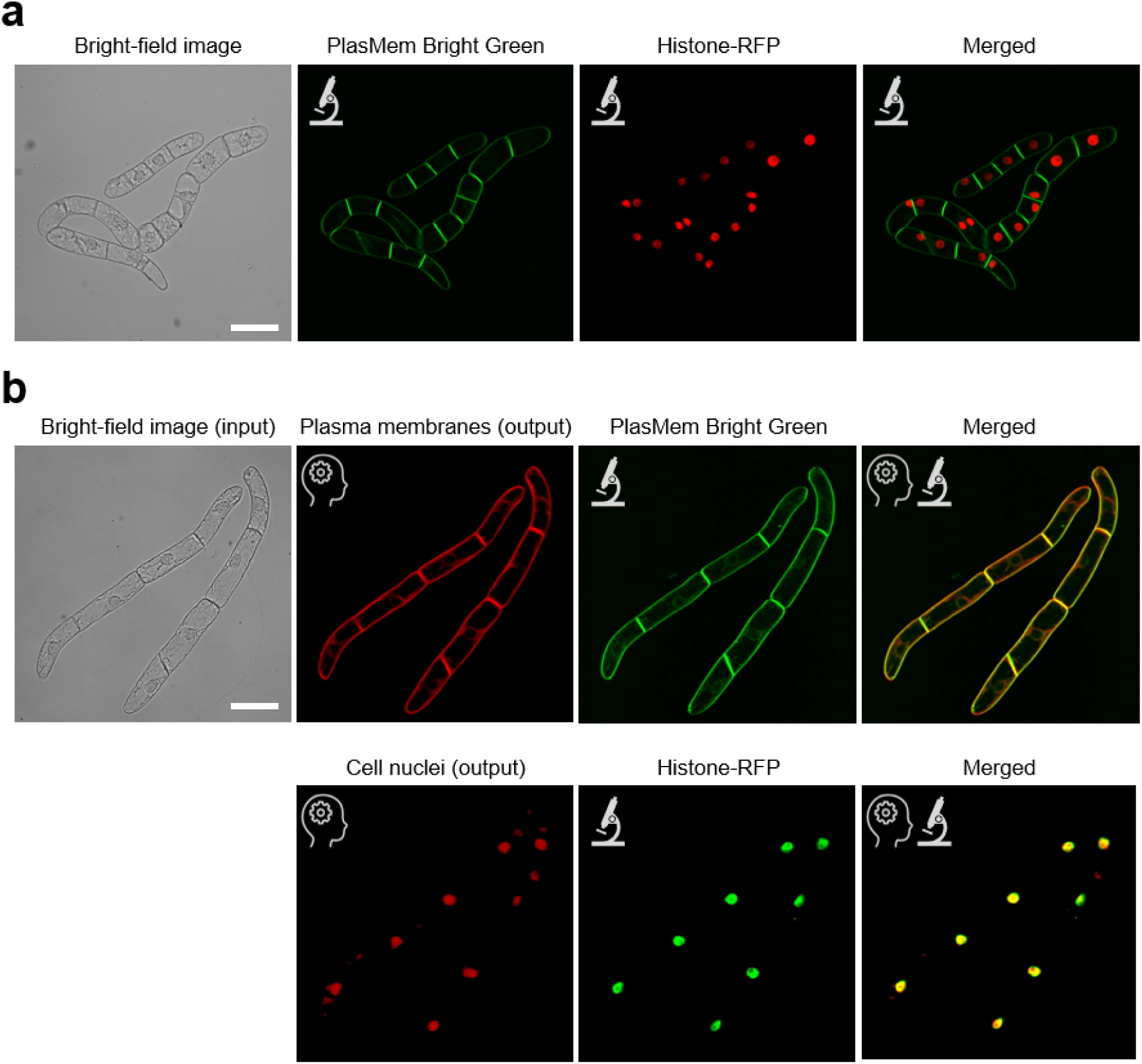
(a) Representative training images. Bright-field microscopy image (far left) is followed by confocal image of the plasma membrane labeled with PlasMem Bright Green (second from left, green), confocal image of the nucleus labeled with Histone-RFP (third from left, red), and merged image of both fluorescent channels (far right). (b) Evaluation of virtual staining. Input bright-field image is shown at top left. Virtual staining outputs for the plasma membrane and cell nuclei are shown in the upper (second from left, red) and lower rows (second from left, red), respectively. For comparison, confocal images of the plasma membrane labeled with PlasMem Bright Green (upper, second from right, green) and the nucleus labeled with Histone-RFP (lower, second from right, green) are shown. Rightmost panels show merged images of virtual staining (red) with actual fluorescence staining (green) for the plasma membrane (upper row) and the nucleus (lower row). Note that the virtual staining images closely match the corresponding physical staining images. AI icon indicates virtual staining images, microscope icon represents actual confocal microscopy images. Scale bars = 100 μm.

**Table 1.** Colocalization analysis between fluorescence-labeled images and virtually stained images. Threshold overlap score (TOS) was used as a colocalization metric, where a TOS value of 1 indicates perfect colocalization between the two images, and lower values reflect decreased colocalization. Mean ± SD. *N*=42.

|  | PlasMem Bright Green | Histone-RFP |
| --- | --- | --- |
| Virtually stained plasma membrane | $0.866 \pm 0.0481$ | $0.266 \pm 0.249$ |
| Virtually stained cell nuclei | $0.465 \pm 0.282$ | $0.798 \pm 0.0760$ |

Additionally, we developed a virtual staining model for vacuoles, a characteristic organelle of plant cells, to explore the potential for multicolor virtual staining. We stained the vacuolar lumen of BY-2 cells with the fluorescent dye BCECF (Kutsuna and Hasezawa 2002) and trained the vacuole virtual staining model using 150 training images. The trained model generated images from bright-field microscopy that closely resembled the BCECF-stained images (Supplementary Fig. S1a). When tested on an independent set of images not used in training, the model achieved a TOS value of 0.783 ± 0.130, confirming its accuracy. Utilizing all three models, we could perform triple virtual staining of the plasma membrane, cell nucleus, and vacuoles from bright-field images (Supplementary Fig. S1b).

Next, to validate the ability of virtual staining to quantitatively assess cellular changes, we performed a quantitative analysis of nuclear size and cell elongation based on virtual staining in BY-2 cells treated with the DNA polymerase inhibitor aphidicolin (Yasuhara and Kitamoto 2014) or the tubulin polymerization inhibitor propyzamide (Katsuta and Shibaoka 1988). Transgenic BY-2 cells, in which microtubules were labeled with YFP-TUB6 and actin filaments were labeled with Lifeact-RFP, were also used (Yasuhara and Kitamoto 2014). These cytoskeletal structures were captured by confocal microscopy, and bright-field images were obtained for virtual staining of the plasma membrane and cell nuclei, respectively. As a result, in addition to the two channels of the fluorescent proteins, the two channels of the virtual stains enabled the visualization of both cytoskeletal and cellular structures simultaneously (Fig. 2a). While physically staining four different cellular structures is technically challenging, these results show that virtual staining can effectively complement fluorescent labeling to expand imaging possibilities. Using these virtual staining images, we semi-automatically segmented the nuclear and cellular regions and measured the nuclear area and the aspect ratio of the cells (Fig. 2b, c). The results show that aphidicolin treatment significantly increased the area of cell nuclei, as well as the aspect ratio of the cells (Fig. 2b, c), and that propyzamide treatment significantly decreased the cell aspect ratio (Fig. 2c). These results were consistent with previous reports that aphidicolin treatment induced nuclear enlargement and cell elongation (Yasuhara and Kitamoto 2014), and that propyzamide treatment suppressed cell elongation (Katsuta and Shibaoka 1988). These results suggest that virtual staining has potential applications in quantitative cell morphometry in response to drug treatments.

**Fig. 2.**
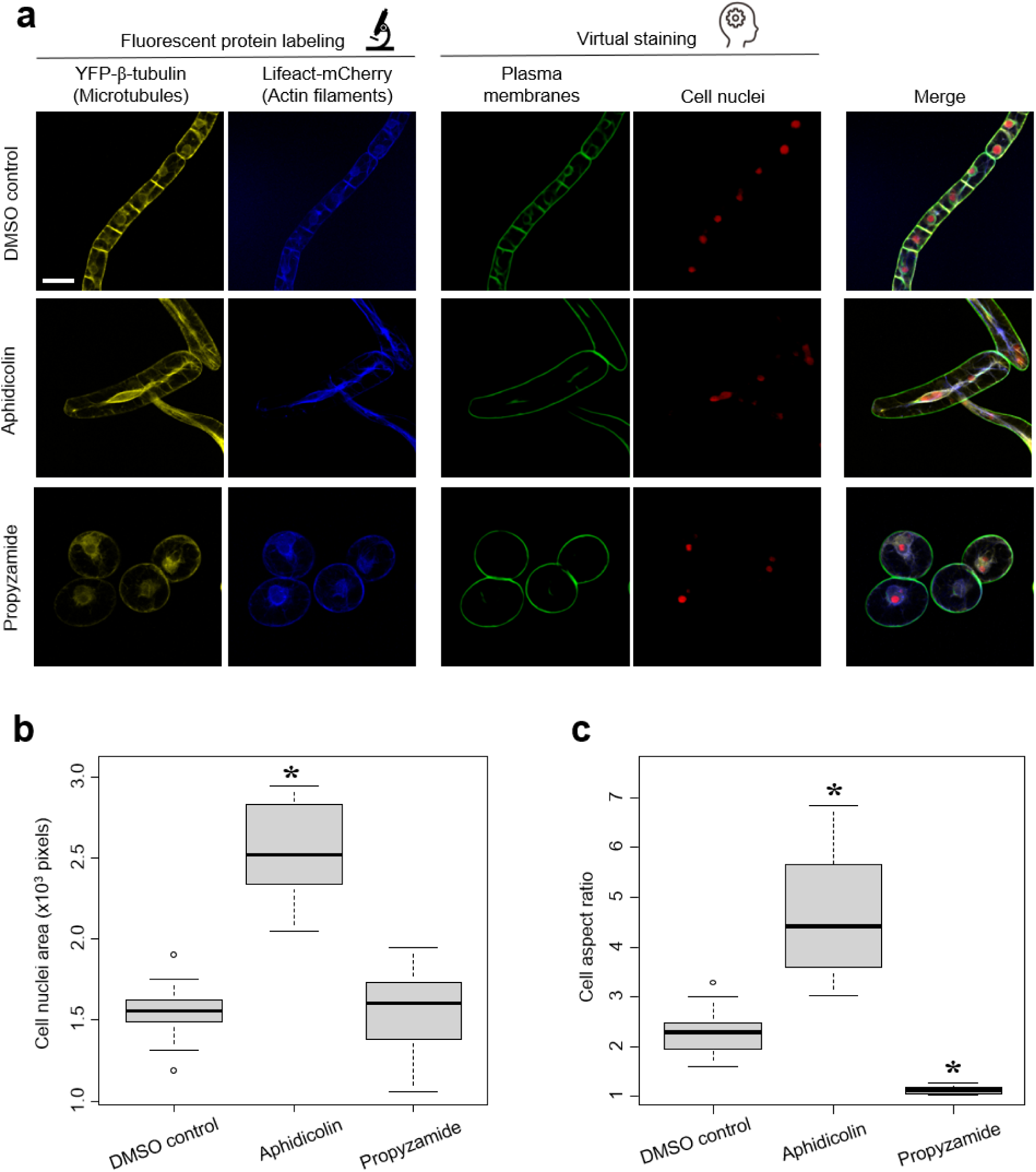
Quadruple staining using virtual staining in tobacco BY-2 cells. (a) Representative images of microtubules labeled with YFP-TUB6 (yellow), actin filaments labeled with Lifeact-RFP (blue), virtually stained plasma membranes (green), and virtually stained cell nuclei (red) in BY-2 cells treated with DMSO control (top), DNA polymerase inhibitor aphidicolin (middle), and microtubule inhibitor propyzamide (bottom). Far right: four-color merged images. Scale bar = 50 μm. (b) Cell nuclei area measured with virtually stained cell nuclei in tobacco BY-2 cells treated with DMSO, aphidicolin, and propyzamide. *P*-values were determined by Mann-Whitney’s U-test (*N* = 15); \**P* < 0.01. (c) Cell aspect ratio measured with virtually stained plasma membrane in tobacco BY-2 cells treated with DMSO, aphidicolin, and propyzamide. Statistically significant differences were determined by Mann-Whitney’s U-test (*N* = 15); \**P* < 0.01.

To further validate the versatility of this method, it was applied to images of *A. thaliana* pavement cells, which have a characteristic puzzle-like morphology and are widely used as a model system in studies on plant cell morphogenesis (Kikukawa et al. 2021a, b). When the virtual staining model for plasma membranes, trained on BY-2 cell images, was applied to the transmitted bright-field images of *A. thaliana* epidermal tissue, output images with visualized plasma membranes were obtained (Fig. 3a, c). Furthermore, semi-automatic segmentation of puzzle-shaped cells was achieved (Legland et al. 2016) (Fig. 3d). To assess the accuracy of virtual staining-based cell segmentation, the cell area and morphological metrics of circularity and solidity (see Methods for their definitions) obtained based on virtual staining images were compared with those obtained by manually tracing from the bright-field microscopy images as ground-truth data (Fig. 3a, b). High correlations (coefficient of determination R^2^ > 0.995) were obtained for all three morphological metrics (Fig. 3e–g). Furthermore, as additional ground-truth data, confocal images of the plasma membrane fluorescent marker GFP-PIP2a were used (Fig. 4). In addition to the bright-field images for virtual staining (Fig. 4a, b), confocal images of GFP-PIP2a were acquired at the same field of view (Fig. 4d). Cells were semi-automatically segmented for each type of image (Fig. 4c, e), and their cell area, circularity, and solidity were measured and compared. High correlations (R^2^ > 0.989) were obtained for all the morphological metrics based on the fluorescent images of the plasma membrane marker and the virtual staining images (Fig. 4f–h). These results show that this virtual staining model for plasma membranes is also valuable for morphometric analysis of pavement cells in *A. thaliana*.

**Fig. 3.**
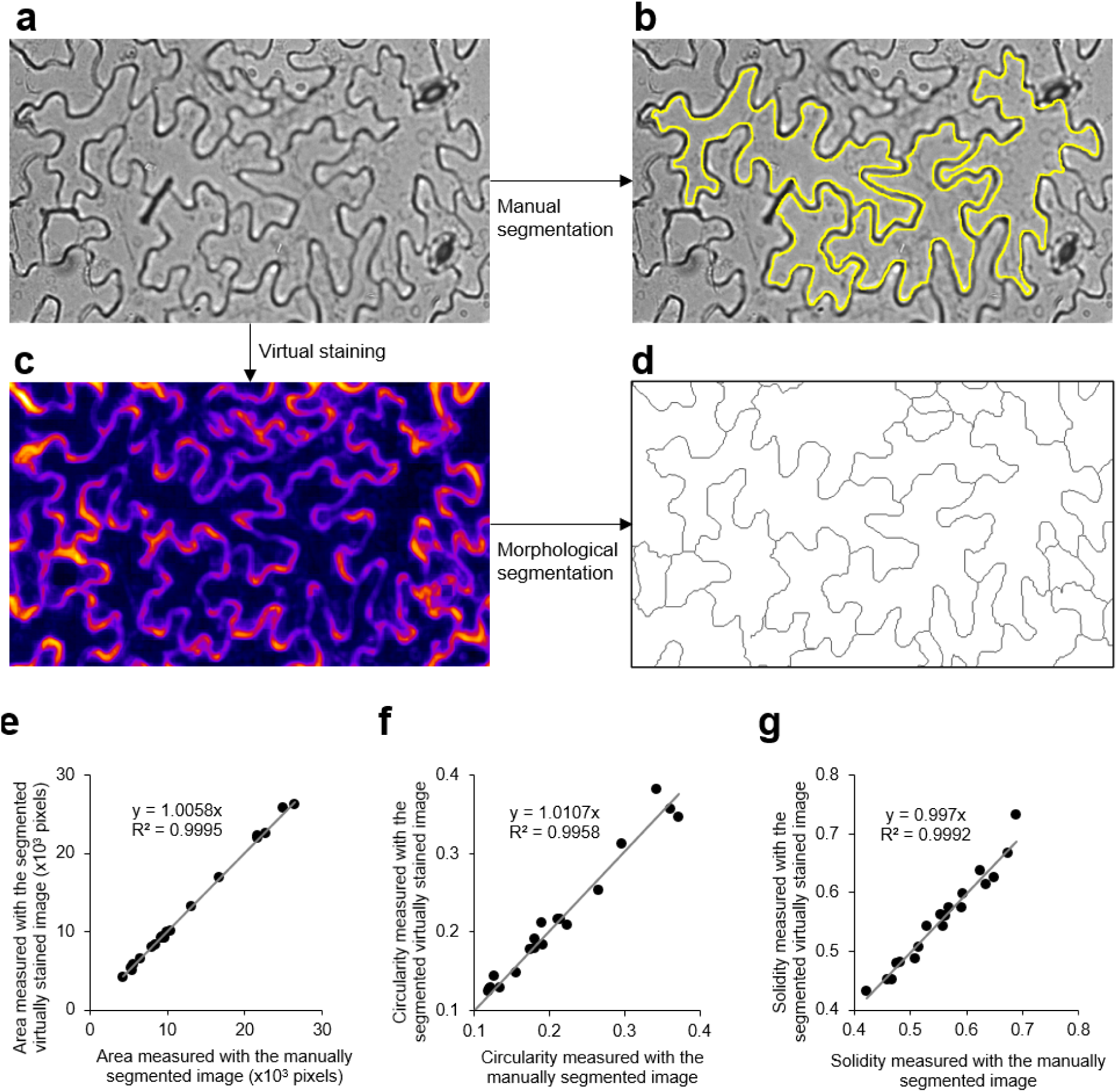
Virtual staining of plasma membranes of *Arabidopsis thaliana* leaf epidermis. (a) Representative transmitted bright-field image of *A. thaliana* leaf epidermis. (b) Manually segmented images of (a). Yellow closed lines indicate contours of pavement cells. (c) Virtually stained image of (a) using virtual staining model trained with bright-field images and confocal images of plasma membranes in tobacco BY-2 cells. (d) Segmented images of (c) using ImageJ plug-in ‘Morphological Segmentation.’ (e–g) Validation of pavement cell segmentation accuracy with measurements of cell area and morphological parameters. Scatter plots of cell area (e), circularity (f), and solidity (g). Relationships between values measured with manually segmented images and virtually stained segmented images are shown. *N* = 20.

**Fig. 4.**
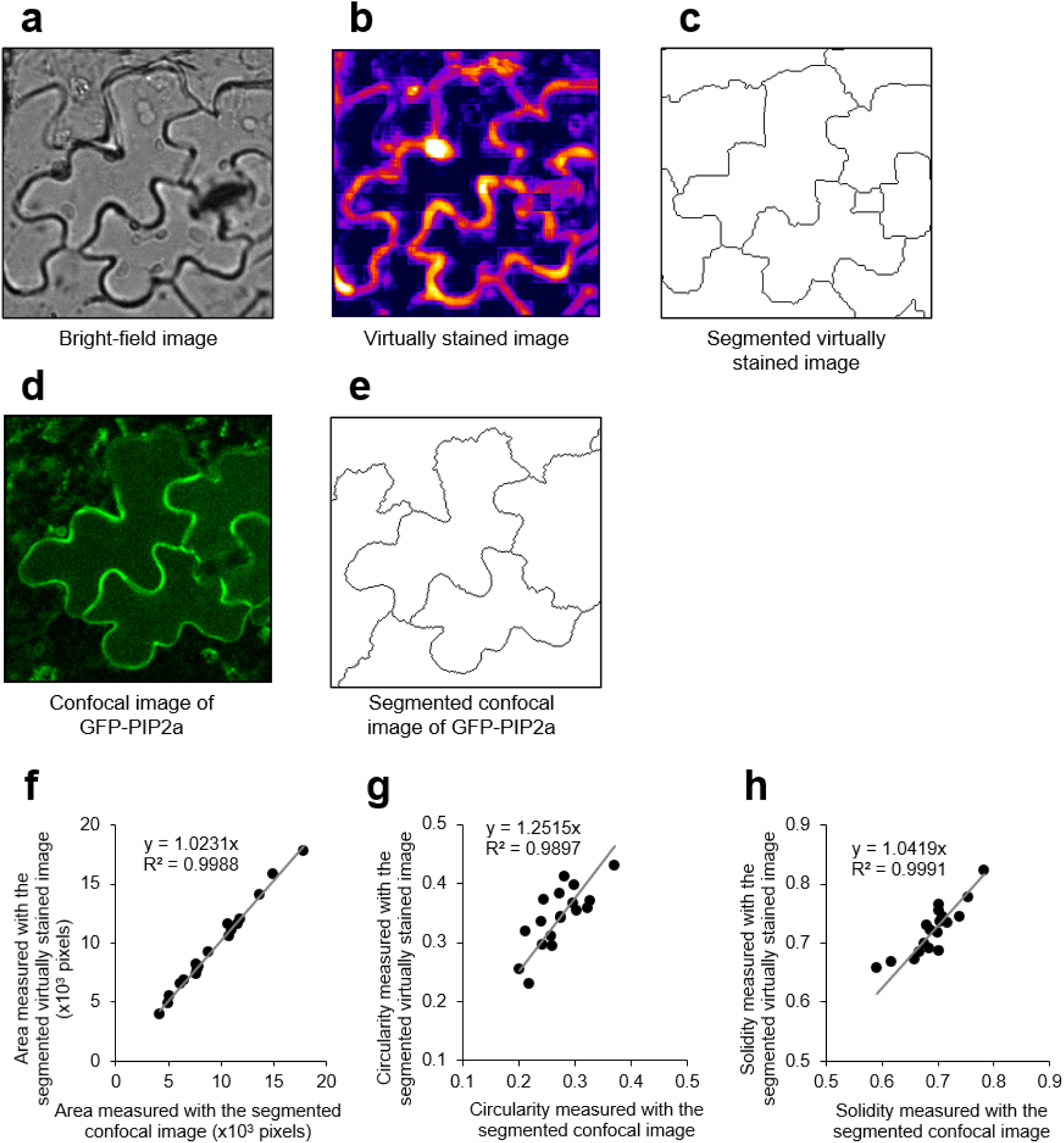
Comparison of confocal images and virtually stained images of plasma membrane markers in *Arabidopsis thaliana* pavement cell morphometry. (a) Representative bright-field image of *A. thaliana* leaf epidermal cells. (b) Virtually stained image of (a) using virtual staining model trained with the tobacco BY-2 cell image data set. (c) Segmented images of (b) using ImageJ plug-in ‘Morphological Segmentation.’ (d) Confocal image of plasma membrane marker GFP-PIP2a within the same field of view as (a). (e) Segmented images of (d) using ImageJ plug-in ‘Morphological Segmentation.’ (f–h) Validation of pavement cell segmentation accuracy with measurements of cell area and morphological parameters. Scatter plots of cell area (f), circularity (g), and solidity (h). Relationships between values measured with segmented confocal images of GFP-PIP2a and segmented virtually stained images are shown. *N* = 18.

To further validate the applicability of this virtual staining-based cell morphometric method for analysis of *A. thaliana* mutants, we used the *bpp125* triple mutant, which exhibits abnormalities in the morphology of pavement cells (Wong et al. 2019; Yoshida et al. 2022). When the virtual staining method for plasma membranes was applied to the transmitted bright-field images of both the wild type and the *bpp125* triple mutant, the plasma membranes were specifically labeled in both genetic backgrounds, and the morphological differences in pavement cells were visualized (Fig. 5a, b). In the *bpp125* triple mutant, the measurements of cell area, circularity, and solidity based on manual tracing showed high correlations with those obtained from virtual staining (R^2^ > 0.998) (Fig. 5c–e). Furthermore, similar to the trend observed in the manual tracing measurements, the circularity and solidity of pavement cells were significantly higher in the *bpp125* triple mutant than in the wild type when measured based on virtual staining (Fig. 5f, g). These results indicate that this method is useful for analyses of genetic mutants, at least in *A. thaliana*.

**Fig. 5.**
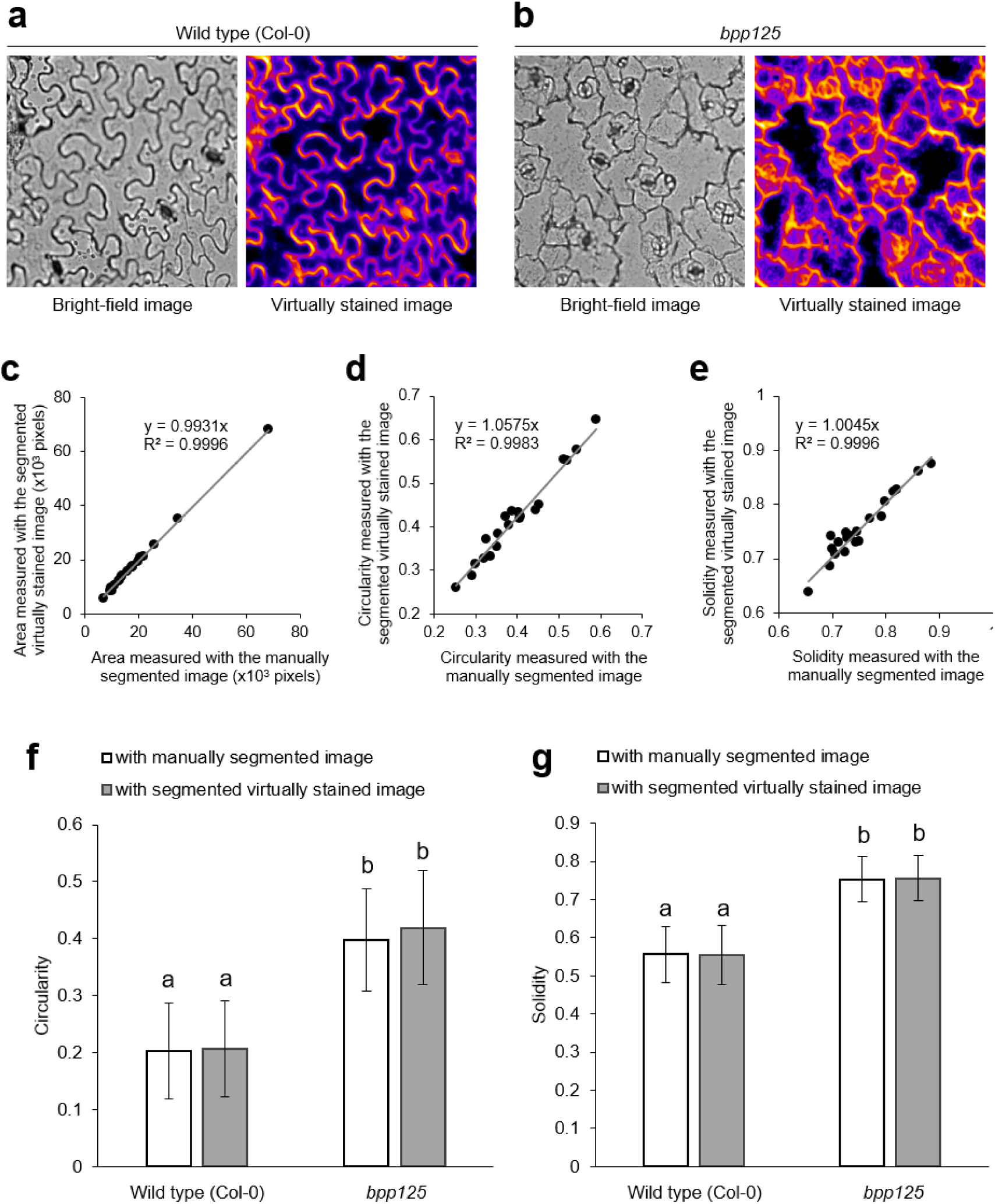
Virtual staining-based phenotyping of *A. thaliana* leaf pavement cell morphology in *bpp125* triple mutant. (a) Representative bright-field image (left) and virtually stained image (right) of *A. thaliana* leaf epidermis of wild type (Col-0). (b) Representative bright-field image (left) and virtually stained image (right) of *A. thaliana* leaf epidermis of *bpp125* triple mutant. (c–e) Validation of pavement cell segmentation accuracy with measurements of cell area and morphological parameters in *bpp125* triple mutant. Scatter plots of cell area (c), circularity (d), and solidity (e). Relationships between values measured with manually segmented images and virtually stained segmented images are shown. *N* = 20. (f, g) Phenotyping of *bpp125* triple mutant. Pavement cell circularity (f) and solidity (g) of wild type (Col-0) (white column) and bpp125 (gray column). Statistically significant differences were detected by Tukey-Kramer test (*N* = 20); \**P* < 0.01.

To evaluate the utility of virtual staining in the analysis of time-lapse image data, we focused on chloroplast movement in *E. densa*, which is well-suited for observing chloroplast movement under bright-field microscopy and has been commonly used in traditional physiological studies (Tazawa et al. 1991). We acquired both bright-field images and autofluorescence images of chloroplasts in the leaflets of *E. densa*. To prevent discrepancies in chloroplast distribution due to their rapid movement caused by cytoplasmic streaming, we treated the samples with BDM, a myosin ATPase inhibitor, to minimize misalignment between the bright-field and autofluorescence images in the training dataset. A total of 109 image sets were used to train the deep learning model, and its accuracy was evaluated on 10 test images that were not included in the training dataset. The trained model generated images that closely resembled the actual chloroplast distribution (Fig. 6a). The TOS value was 0.828 ± 0.0494 (*N*=10), comparable to the accuracy achieved in the virtual staining of plasma membranes and cell nuclei in BY-2 cells (Table 1). To further assess the applicability of virtual staining for evaluating chloroplast movement, we first manually tracked the chloroplast centroid coordinates in bright-field time-lapse images captured every 0.5 s to calculate the speed of chloroplast movement between frames (Fig. 6b). In parallel, we segmented chloroplasts from the virtually stained images using Otsu’s method, automatically determining chloroplast movement speed based on centroid positions (Fig. 6c, d). The manually measured and virtual staining-based automated chloroplast movement speeds showed a strong correlation (Fig. 6e, R² = 0.985). These results show that virtual staining is also effective for analyzing chloroplast movement in time-lapse imaging.

**Fig. 6.**
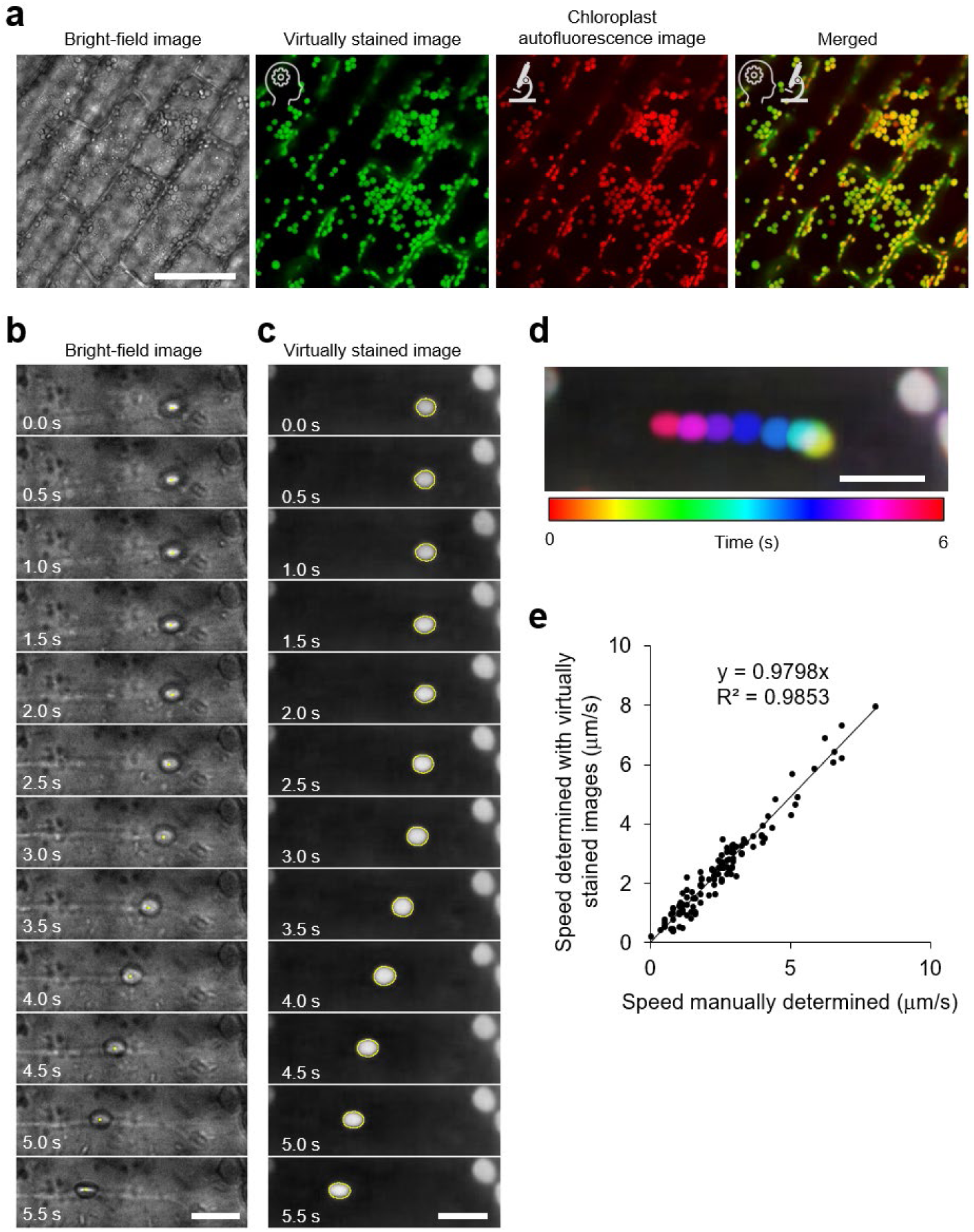
Virtual staining-based evaluation of chloroplast movement speed in *Egeria densa*. (a) Virtual staining of chloroplasts in *E. densa*. Bright-field image (far left) is followed by virtually stained chloroplasts (second from left, green), confocal autofluorescence image of chloroplasts (second from right, red), and merged image of both virtual staining and autofluorescence (far right). AI icon indicates virtual staining images, microscope icon represents actual confocal microscopy images. Scale bar = 100 μm. (b) Chloroplast movement as observed in bright-field images. Chloroplast centroids (yellow dots) were manually determined to calculate movement speed between frames. Scale bar = 10 μm. (c) Chloroplast movement based on virtually stained images. Chloroplasts were segmented using thresholding (yellow outlines), and movement speed was calculated from the centroids. Scale bar = 10 μm. (d) Visualization of chloroplast movement using a temporal color code. Overlaid images show chloroplast movement across different time frames, each represented by a distinct color. Scale bar = 10 μm. (e) Evaluation of chloroplast movement speed determined using virtual staining. Speed calculated from manually tracked bright-field images showed a high correlation (R² = 0.9853) with that determined from virtually stained images.

To evaluate the utility of virtual staining in the automatic recognition of cell states, we captured 1,200 bright-field images each of live BY-2 cells treated with 0.1% DMSO as a control and of dead cells treated with 10 μM enoxolone, a compound we identified as a cell death inducer in these cells after 24 hours, using the automated imaging system BioTek Cytation1. When virtual staining of the plasma membrane was applied to these images, live cells displayed clear and intact cell contours, while dead cells exhibited shrunken and collapsed internal cell membranes (Fig. 7a). We then performed cell viability classification using machine learning on both bright-field and virtually stained images. GLCM features, a commonly used texture descriptor (Haralick 1979), were extracted from 900 images of live cells and 900 images of dead cells (in total, 1,800 images). These texture features were used to train a random forest classifier, and classification accuracy was assessed using the remaining 300 images of each cell status (in total, 600 test images). Ten classifiers were generated with different random seeds to validate classification accuracy. The results showed that the classification accuracy using bright-field images was 76.2% ± 0.277%, while that using virtually stained images was significantly higher at 87.9% ± 0.309% (Fig. 7b, Tables 2 and 3). This finding shows that virtual staining offers superior accuracy in determining cell viability.

**Fig. 7.**
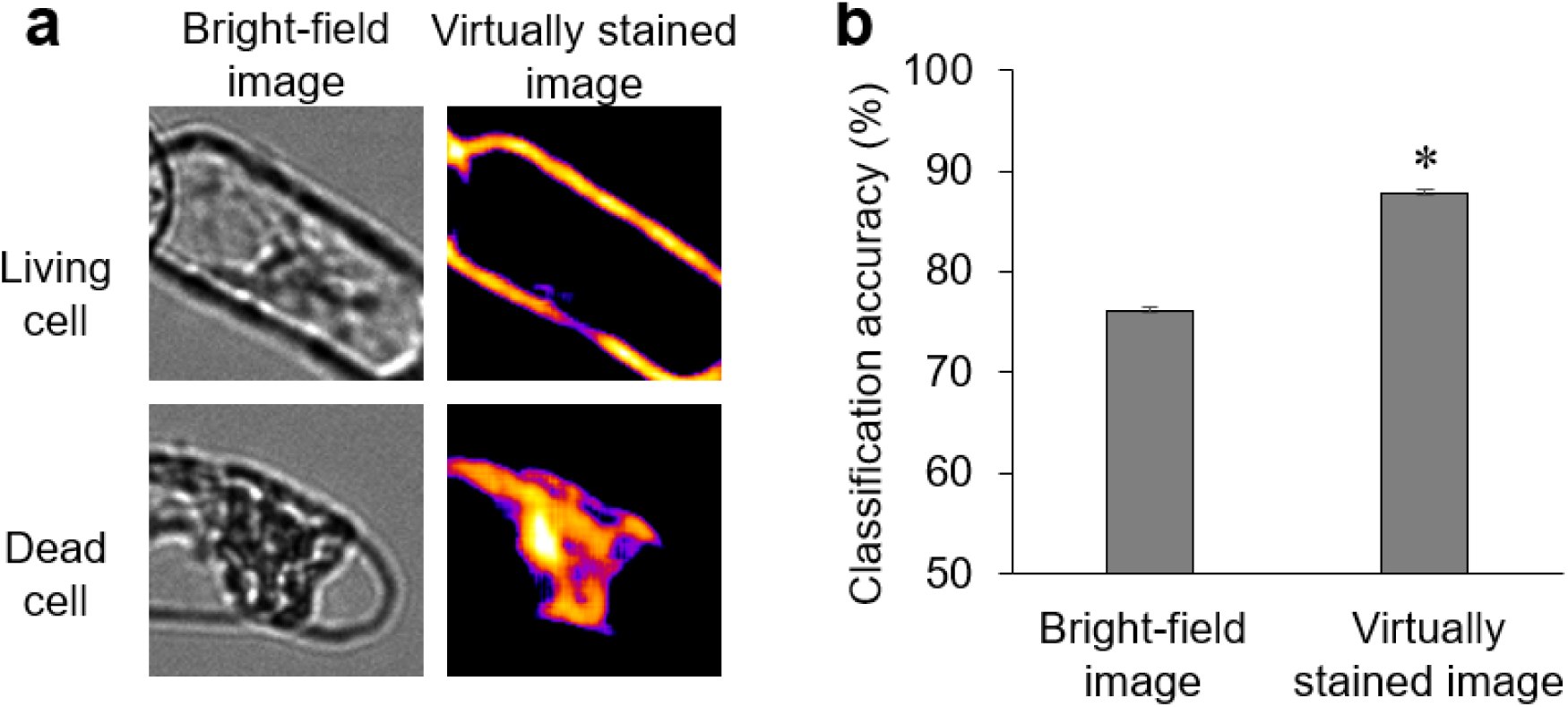
Enhanced accuracy of cell viability classification with virtual staining in tobacco BY-2 cells. (a) Representative images of living (top) and dead (bottom) tobacco BY-2 cells. Bright-field images (left) display general morphology, while virtually stained images (right) highlight the plasma membrane, allowing for clearer distinction between living and dead cells. (b) Classification accuracy of cell viability based on bright-field and virtually stained images. Random forest classification was applied using GLCM-based texture features. Classification accuracy was significantly better when using virtually stained images than when using bright-field images (p < 0.01, U-test). Confusion matrices for bright-field and virtually stained images are provided in Tables 2 and 3.

**Table 2.** Confusion matrix for cell viability classification based on bright-field images using random forest classification. Classification performance for distinguishing between living and dead cells is shown. The matrix shows the number of true living and dead cells against their predicted classifications.

|  | Predicted living cells | Predicted dead cells |
| --- | --- | --- |
| Living cells | 221 | 79 |
| Dead cells | 65 | 235 |

**Table 3.** Confusion matrix for cell viability classification based on virtually stained images using random forest classification. Classification performance for distinguishing between living and dead cells using virtually stained images is shown. The matrix presents the number of true living and dead cells against their predicted classifications.

|  | Predicted living cells | Predicted dead cells |
| --- | --- | --- |
| Living cells | 263 | 37 |
| Dead cells | 34 | 266 |

## Discussion

In this study, we investigated whether virtual staining based on deep learning, which is advancing mainly in the biomedical field, was also applicable to plant cells. In animal cells, virtual staining has been reported to achieve high accuracy in detecting structures that are easily visible in bright-field images, such as lipid droplets, cell nuclei, and nucleoli (Wieslander et al. 2021; Cross-Zamirski et al. 2022). As shown in this study, virtual staining successfully distinguished not only cellular structures common to both plant and animal cells, such as plasma membranes and cell nuclei, but also plant-specific organelles like vacuoles and chloroplasts (Supplementary Fig. S1, Fig. 6). These results show that the method can be adapted for use in plant biology. To quantitatively assess the effectiveness and accuracy of virtual staining in plant cells, we utilized the TOS (Stauffer et al. 2018; Hotta et al. 2022). The high TOS values observed for these cellular structures indicate that virtual staining can reliably replicate plant cell structures (Table 1).

While virtual staining based on bright-field microscopic images has promising applications in plant cell research, there are certain technical limitations to consider. For instance, in this study, we used plant samples consisting of a single layer of cells, such as tobacco BY-2 cells and the epidermal tissue of *A. thaliana* leaves. However, tissues or organs composed of multiple layers of relatively small cells, such as *A. thaliana* roots, may not be suitable for virtual staining due to the difficulty in discerning individual cell structures with bright-field microscopy. Additionally, unlike confocal microscopy, virtual staining based on bright-field images is inherently limited to two-dimensional information, and cannot readily provide three-dimensional structural data, which could be critical for more complex tissue analysis (Higaki and Mizuno 2020).

Despite these limitations, virtual staining has several advantages that make it a valuable complement to traditional fluorescent labeling techniques. In this study, we were able to detect nuclear enlargement and cell elongation induced by aphidicolin, as well as the inhibition of cell elongation by propyzamide, in BY-2 cells (Fig. 2). Furthermore, in the pavement cells of *A. thaliana* leaves, the results of cell morphometry based on virtual staining closely matched those obtained from manual tracing of bright-field microscopy images (Figs. 3 and 5) or semi-automatic segmentation based on the plasma membrane marker GFP-PIP2a (Fig. 4). The ability to virtually stain cellular structures, such as the plasma membrane and cell nuclei, provides a flexible approach that can alleviate the need for physical fluorescent labeling. This flexibility enables researchers to allocate fluorescent resources to other cellular components. For example, virtual staining of the plasma membrane and nucleus can free up fluorescent channels for multiple cytoskeletal markers, potentially facilitating more detailed investigations of cellular organization and dynamics (Fig. 2a). This approach simplifies experimental design, particularly in multicolor labeling setups that involve complex cellular events. Our analysis of chloroplast movement in *E. densa* further demonstrates the versatility of virtual staining for dynamic imaging applications (Fig. 6). Virtual staining enabled automated tracking of chloroplasts over time-lapse sequences, and the results showed a strong correlation with those obtained by manual tracking (R² = 0.985) (Fig. 6e). This approach allows for efficient, label-free analysis of intracellular movement without the need for fluorescent dyes and the use of excitation lasers, which can lead to photobleaching and potential cytotoxicity. The high accuracy of virtual staining in tracking chloroplast movement suggests that this method could be beneficial for studying other plant-specific dynamic processes, such as cytoplasmic streaming and organelle transport, particularly in time-lapse imaging setups. Furthermore, our classification of live and dead BY-2 cells highlights the potential of virtual staining in non-segmentation-based analyses (Fig. 7). Using virtually stained images, we achieved a significantly higher classification accuracy for live and dead cells compared with that using bright-field images alone, with improvement of accuracy from 76.2% to 87.9% (Fig. 7b, Tables 2 and 3). This result shows that virtual staining can enhance the ability to differentiate between living and dead cells without the need for fluorescent viability dyes. Unlike segmentation-based methods, this approach leverages texture-based machine learning techniques, such as GLCM feature extraction, to enable effective and automated classification of cell state. Consequently, virtual staining offers a rapid, label-free alternative for viability assays in plant cells, with potential applications in cytotoxicity testing and other high-throughput screening protocols.

As noted, virtual staining is not intended to replace fluorescence imaging entirely, as fluorescence imaging remains an essential tool in cell biology. However, by effectively integrating virtual staining with fluorescence imaging, researchers can extend the possibilities for imaging experiments in plant cell biology. Notably, virtual staining offers advantages such as reduced phototoxicity and the elimination of the need for fluorescent dyes, which will be beneficial for time-lapse imaging studies and phenotypic analyses of mutants. Therefore, our findings demonstrate that virtual staining based on deep learning is a powerful tool for label-free visualization and precise morphometry of plant cell structures. By complementing traditional techniques, virtual staining can enhance the overall imaging capabilities in plant cell biology research.

In conclusion, the results of this study show that virtual staining based on deep learning is a valuable tool for label-free visualization and precise morphometry of plant cell structures. While this method does not replace fluorescence imaging, it offers a complementary approach that overcomes some of the limitations associated with traditional imaging techniques, such as phototoxicity and the need for fluorescent labeling. Our results suggest that virtual staining could significantly enhance plant cell biology analyses, especially in studies involving time-lapse imaging and phenotypic analyses of mutants.

## Author Contributions

Takumi Higaki contributed to the study conception and design. Material preparation, data collection and analysis were performed by Manami Ichita, Haruna Yamamichi, and Takumi Higaki. The first draft of the manuscript was written by Takumi Higaki and Manami Ichita commented on previous versions of the manuscript. All authors read and approved the final manuscript.

## Data Availability

The data pertaining to this article will be shared on reasonable request to the corresponding author. The training and test image sets for BY-2 cells and *E. densa*, which are publicly accessible on figshare under the CC BY 4.0 license, include images of wild-type tobacco BY-2 cells stained with BCECF for vacuolar lumen visualization (https://doi.org/10.6084/m9.figshare.27247629.v1), transgenic tobacco BY-2 cells with Histone-RFP and PlasMem Bright Green staining (https://doi.org/10.6084/m9.figshare.27247638.v1), and *E. densa* chloroplast autofluorescence images (https://doi.org/10.6084/m9.figshare.27247620.v1).

## Acknowledgements

We thank Ms. Hitomi Okada (Kumamoto University), Ms. Remi Kawakami (Kumamoto University), and Ms. Koko Ishida (Kumamoto University) for their assistance with plant maintenance and image acquisition. We also thank Jennifer Smith, PhD, from Edanz Group (https://jp.edanz.com/), for editing a draft of this manuscript.

## Supplementary Figure

**Supplementary Fig. S1.**
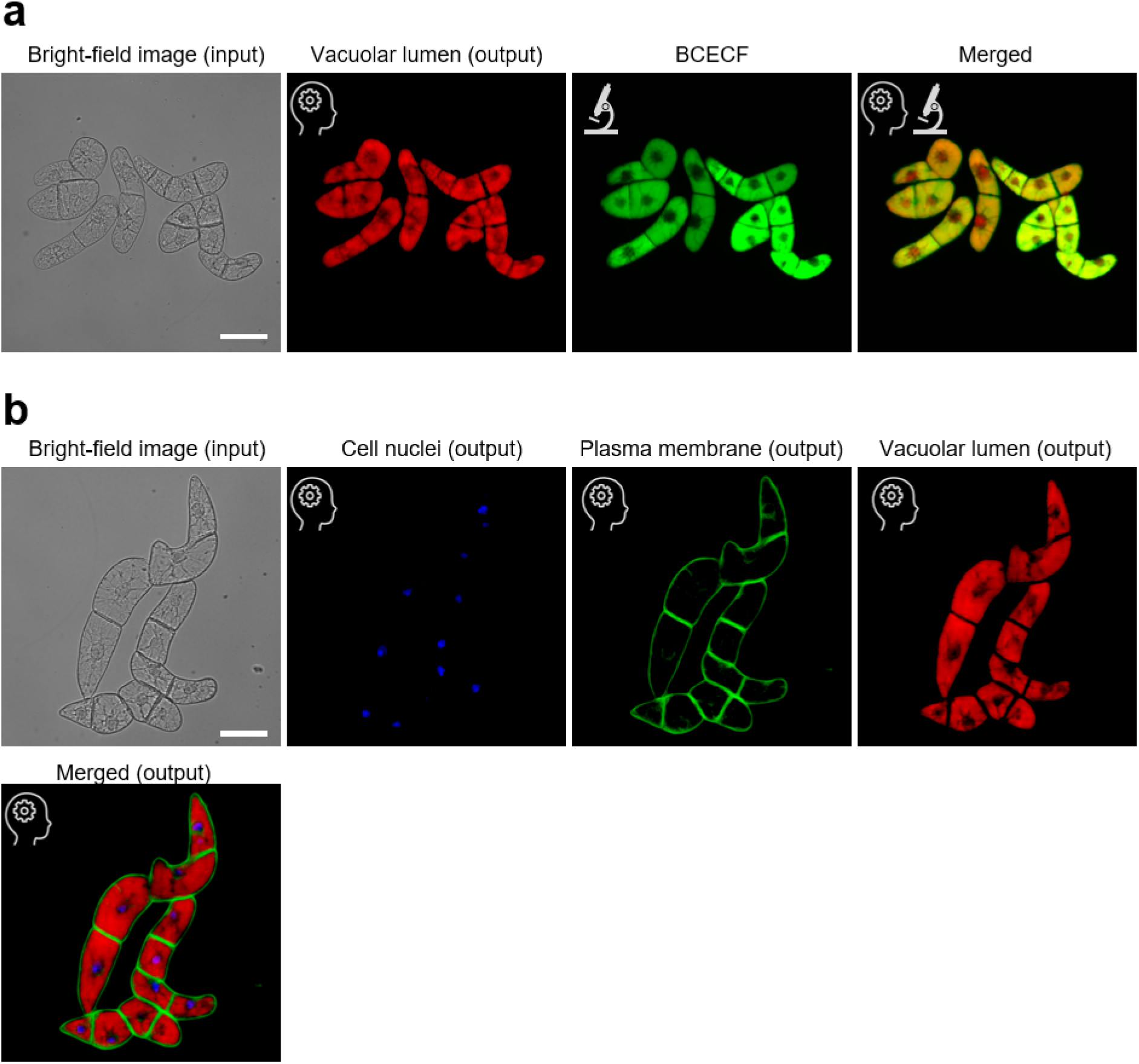
Virtual staining of vacuoles in tobacco BY-2 cells. (a) Evaluation of virtual staining. Input bright-field image (far left) is followed by virtual staining output of the vacuolar lumen (second from left, red), confocal image of the vacuolar lumen labeled with the fluorescent dye BCECF (second from right, green), and merged image of virtual staining and fluorescence staining (far right). Virtually stained images closely align with corresponding confocal images. (b) Triple virtual staining of BY-2 cells. Bright-field image (far left) is followed by virtual staining outputs for cell nuclei (blue), plasma membrane (green), and vacuolar lumen (red). Lower panel displays a merged image of these three virtual stains. AI icon indicates virtual staining images, microscope icon denotes actual confocal microscopy images. Scale bars = 100 μm.

